# Sex differences in auditory function of the desert locust

**DOI:** 10.1101/2024.11.13.622961

**Authors:** Austin T Tom, Thomas L Christian, Warren Ben

## Abstract

Age-related auditory decline manifests across the animal kingdom, from humans and mice to zebrafish and insects. Sex differences in auditory decline are established for humans, but there is now evidence in mice and even zebrafish. Here, we found sex differences in auditory decline in an insect, the Desert Locust and investigated its biological basis. We profiled gene expression in a dedicated auditory organ, Müller’s organ to understand the genetic underpinning of sex differences and measured sound-evoked transduction currents and electrophysiological properties of auditory neurons to quantify auditory decline. We analysed gene expression in Müller’s organ of young locusts where sex differences were absent and in older, noise-exposed locusts where sex differences were maximal. The largest differences in gene expression between the sexes was between young and stressed (aged and noise-exposed) auditory organs. We found sex-specific genes and gene ontology terms for juvenile hormone (JH) and sex-specific estrogen-related steroids. We hypothesise that sex differences in auditory decline are due to differences in hormones which then affect metabolic processes in and mitochondria.

**Highlights:**

- Female Desert Locusts have less auditory decline than males.
- Female’s auditory organs maintain better metabolism under stressed conditions.
- Sex differences in gene expression are maximal during aging and noise-exposure.
- Sex differences are minimal between the sexes for young and stressed auditory organs.

## Introduction

Hearing loss in an inevitable consequence of aging. But for females hearing loss is less for higher frequencies >4 kHz (Spoor, 1967; Corso, 1963), whereas males hearing is better for low frequencies <500 Hz hearing. Historically these sex differences were assumed to be due to sex differences in noise exposure in the male-dominated noisy workplaces of the mid-to late 20^th^ Century (Gates et al., 1990). Later, more numerous studies, that accounted for noise, and that were based on modern cohorts in quieter work environments with more equal gender balance revealed conclusively that females have less auditory decline, independent of environmental factors (Wang et al., 2021; Homans et al, 2017; Gablenz & Holube, 2016; Allen and Eddins, 2010; Agrawal et al., 2008; Helzner et al., 2005; Pearson et al., 1995, Cruickshanks et al., 1998). Once it was established that something inherent in the biology of females led to less hearing loss researchers started to search for its physiological basis.

One of the strong candidates are female-specific sex hormones such as progesterone and estrogen (Nolan, 2020). Indeed, a range of hearing measures correlate with the menstrual cycle in humans (Al-Mana, et al., 2010). In the popular hearing model, *Mus musculus*, there are conflicting findings of sex differences, with some finding a difference (Milon et al., 2018) and other, larger powered studies, not (Ingham et al., 2019, Fig. S2). However, these conflicting accounts could be due to an overriding effect of the menstrual cycle, that averages out sex differences. In the vertebrate model *Danio reiro*, pure-tone noise exposure led to consistently larger threshold shifts for males that were postulated due to higher levels of damage represented by the higher levels of cortisol in males (Han el., 2022). It should be noted that the effect size of the sex differences found in humans is small and typically no larger than 10 dB SPL, and thus only found with a large enough sample size (Nolan, 2020). Sex differences in hormones by themselves do not explain the physiological mechanism of hearing differences; the answer relies on identifying the cellular pathways, processes and organelles on which the hormones act. Strong candidates include metabolic pathways in mitochondria. Not only do mitochondria have receptors for hormones (Yang et al., 2004), but the once (pre-eukaryotic) symbiotic α-protobacteria hold an ancient evolutionary reason that could explain male-biased hearing loss.

Almost all mitochondria in eukaryotes are inherited through the maternal germline. This means that natural selection acts only on the mitochondria in females and that mutations in mitochondria that might be deleterious for males are less so for females; This theory is also known as the “Mother’s curse”. Mitochondria are integral to eukaryote cellular function, so much so that most of the ∼1,600 mitochondrial genes are now coded in the nuclear DNA, leaving only 37 still on the circular DNA of the mitochondria itself (Gray, 1992; Johnston & Williams, 2016). This results in extensive mitochondria-nuclear cross-talk necessary to coordinate proteins necessary for mitochondrial function and also to signal the energy demands of the cell. This mitochondria-nuclear communication is hypothesised to work better in females. We hypothesise that sex-differences in mitochondrial function only become apparent under stress induced by age or use, which could explain sex differences in hearing only in older cohorts.

Auditory organs themselves have greater metabolic demands than other cell types. For instance, the receptor cells are bathed in a haemolymph (scala media in mammals) that creates an electrochemical gradient of +140 mV, twice as high compared to neurons. Cells dedicated to establishing this electrochemical gradient are thought to be the most metabolically active and therefore the target of metabolic-based insults (Schuknecht and Gacek, 1993; Warren et al., 2020). In insects, a metabolically demanding scolopale cell maintains a specialised receptor lymph (analogous to the scala media) of the sensory cilium. In addition, higher frequency-parts of auditory organs maintain a metabolically-demanding higher electrochemical gradient, both in mammals and insects (Salt et al., 1989; Sterkers et al., 1984; Manuela Nowotny personal communication), possibly explaining why high frequency hearing is better in females. Another compounding explanation for high-frequency hearing loss is the higher demands for Ca^2+^ buffering, in high frequency regions of the cochlea, where the smaller receptor cells have less inherent buffering ability (Fettiplace & Nam, 2019).

If the mitochondria hypothesis is correct, we would expect to see sex-biased mitochondrial function across eukaryotic auditory organs, not just in mammals. There is some evidence for this in the auditory organs of zebrafish (Han et a., 2022), but, in theory, these findings should extend further across the animal kingdom. We wanted to investigate if sexual differences in auditory decline exist in the Desert Locust, *Schistocerca gregaria*, and if so to understand their physiological basis. To do this we measured sound-evoked transduction currents and electrophysiological properties from individual auditory neurons as a measure of auditory function and then measured gene expression, to understand the physiological basis of any differences. In order to maximise the chances of finding sexual differences and identifying their cause, we used a young cohort and a second aged and noise-exposed cohort.

## Methodology

### Locust Husbandry

*Schistocerca gregaria* (Mixed sex) were raised in crowded conditions 150–250 in 60 cm^3^ cages in their fast-aging gregarious state, where they can live up to two months. This contrasts with their isolated solitarious state where they live for up to nine months. *S. gregaria* where fed ab libitum fresh wheat (grown in house) and milled bran. The founding progeny of the Leicester Labs strain were solitary copulating adults collected at Akjoujt station ∼250 km North East from Nouakchott, Mauritania in May 2015.

### Noise exposure, aging and acoustic stimulation

No anaesthesia was used for experiments with locusts. Experiments were performed on four cohorts of the locusts comprising both sexes (four groups). The young group were 10 days post their last moult, the aged and noise-exposed group (here after referred to as the stressed group) were 28 days post their last moult (**Figure 1**). The wings of all locusts (young and stressed) were cut off at their base to increase noise exposure of the conditioning tone to their tympanal ears, which are otherwise covered by their wings. Between ten and twenty locusts, for both the control group and the stressed group, were placed in a cylindrical wire mesh cage (8 cm diameter, 11 cm height). Both cages were placed directly under a speaker (Visaton FR 10 HM 4 OHM, RS Components Ltd). For the stressed group only, the speaker was driven by a function generator (Thurlby Thandar Instruments TG550, RS Components Ltd) and a sound amplifier (Monacor PA-702, Insight Direct Ltd) to produce a 3 kHz tone at 120 dB SPL (Sound Pressure Level), measured at the top of the cage where locusts tended to accumulate. Throughout the paper we refer to noise that the locusts are exposed to as a 3 kHz 120 dB SPL pure tone. This tone was played continuously for 12 h overnight (21:00-09:00) for the stressed group during their natural darkness period. The noise-exposure was repeated for the stressed group very three days a total of six times. The stressed group were experimented on (ears extracted for RNA extraction or ear excised for patch-clamp electrophysiology) between 48-60 hours after the last noise exposure (**Figure 1A**). The control group was housed in an identical cage with a silent speaker for 12 h. All recordings were performed within a 12 h window during the day. Sound Pressure Levels (SPLs) were measured with a microphone (Pre-03 Audiomatica, DBS Audio) and amplifier (Clio Pre 01 preamp, DBS Audio). The microphone was calibrated with a B&K Sound Level Calibrator (CAL73, Mouser Electronics). For patch-clamp recordings, the locust ear was stimulated with the same speaker and amplifier as above with a 3 kHz pure tone duration of 0.5 s Tones were played three times for each locust at each SPL and the average response taken for each SPL. For intracellular recordings from individual auditory neurons the speaker was driven by a custom-made amplifier controlled by an EPC10-USB patch-clamp amplifier (HEKA-Elektronik) controlled by the program Patchmaster (version 2 × 90.2, HEKA-Elektronik) running under Microsoft Windows (version 10).

**Figure 1.**
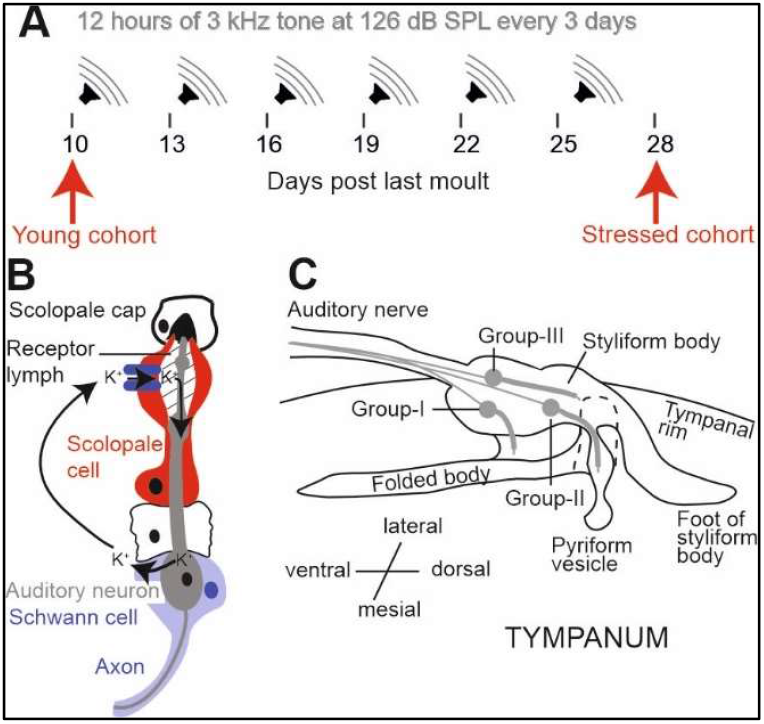
Workflow of experiments and anatomy of Müller’s organ. **A**. Experimental showing when the cohorts of locusts were experimented. Young locusts were taken at 10 days post last moult (red arrow) and stressed locusts were taken at 28 days post last moult and after six overnight noise exposures (red arrow). **B**. Diagram of the sensory unit (scolopidia) of the insect auditory system. The red scolopale cell recycles K^+^ and is assumed to be highly metabolically active. **C**. Diagram of Müller’s organ with each group of auditory neurons represented by one neuron in red. Müller’s organ is attached to the tympanum at the folded body (group-I), foot of styliform body (group-III) and pyriform vesicle (Group-II).

### Dissection of Müller’s organ and isolation of Group-III auditory neurons

Whole cell patch clamp recordings were performed on group-III auditory neurons because they form the majority of auditory neurons of Müller’s organ (∼46 out of ∼80) (Jacobs et al., 1999), they are the most sensitive auditory neurons of Müller’s organ (Römer, 1976) and are broadly tuned to the 3 kHz we used for noise-exposure (Warren and Matheson, 2018). For intracellular patch-clamp recordings from individual auditory neurons the abdominal ear, including Müller’s Organ attached to the internal side of the tympanum (**Figure 1 B, C**), was excised from the first abdominal segment, by cutting around the small rim of cuticle surrounding the tympanum with a fine razor blade. Trachea and the auditory nerve (Nerve 6) were cut with fine scissors (5200-00, Fine Science Tools), and the trachea and connective tissue removed with fine forceps. This preparation allowed perfusion of saline to the internal side of the tympanum, necessary for water-immersion optics for visualizing Müller’s Organ and the auditory neurons to be patch-clamped, and concurrent acoustic stimulation to the dry external side of the tympanum. The inside of the tympanum including Müller’s Organ was constantly perfused in extracellular saline. Dissection, protease and recordings took ∼60 min for each locust ear.

To expose Group-III auditory neurons for patch-clamp recordings, a solution of collagenase (0.5 mg/mL) and hyaluronidase (0.5 mg/mL) (C5138, H2126, Sigma Aldrich) in extracellular saline was applied onto the medial-dorsal border of Müller’s Organ through a wide (12 μm) patch pipette to digest the capsule enclosing Müller’s Organ and the Schwann cells surrounding the auditory neurons. Gentle suction was used through the same pipette to remove the softened material and expose the membrane of Group-III auditory neurons. The somata were visualized with a Cerna mini microscope (SFM2, Thor Labs), equipped with infrared LED light source and a water immersion objective (NIR Apo, 40×, 0.8 numerical aperture, 3.5 mm working distance, Nikon) and multiple other custom modifications.

### Electrophysiological recordings and isolation of the transduction current

Electrodes with tip resistances between 3 and 4 MΩ were fashioned from borosilicate class (0.86 mm inner diameter, 1.5 mm outer diameter; GB150-8P, Science Products GmbH) with a vertical pipette puller (PC-100, Narishige). Recording pipettes were filled with intracellular saline containing the following (in mM): 170 K-aspartate, 4 NaCl, 2 MgCl2, 1 CaCl2, 10 HEPES, 10 EGTA. 20 TEACl. Intracellular tetraethylammonium chloride (TEA) was used to block K+ channels necessary for isolation the transduction. To further isolate and increase the transduction current we also blocked voltage-gated sodium channels with 90 nM Tetrodotoxin (TTX) in the extracellular saline. During experiments, Müller’s Organs were perfused constantly with extracellular saline containing the following in mM: 185 NaCl, 10 KCl, 2 MgCl2, 2 CaCl2, 10 HEPES, 10 Trehalose, 10 Glucose. The saline was adjusted to pH 7.2 using NaOH. The osmolality of the intracellular and extracellular salines’ were 417 and 432 mOsm, respectively.

Whole-cell voltage-clamp recordings were performed with an EPC10-USB Double patch-clamp amplifier (HEKA-Elektronik) controlled by the program Patchmaster (version 2 × 90.2, HEKA-Elektronik) running under Microsoft Windows (version 10). Electrophysiological data were sampled at 50 kHz. Voltage-clamp recordings were low-pass filtered at 2.9 kHz with a four-pole Bessel filter. Compensation of the offset potential were performed using the “automatic mode” of the EPC10 amplifier and the capacitive current was compensated manually. The calculated liquid junction potential between the intracellular and extracellular solutions was also compensated (15.6 mV; calculated with Patcher’s-PowerTools plug-in from www3.mpibpc.mpg.de/groups/neher/index.php?page=software). Series resistance was compensated at 77% with a time constant of 100 μs.

### RNA extraction of Müller’s organs and RNA sequencing

Müller’s organ were extracted by holding live locusts to expose the tympanum in the 1^st^ abdominal segment. Fine forceps pierced through the tympanum either side of Müller’s organ before being closed to grasp Müller’s organ. Müller’s organ was then wiped onto a pestle and placed into an Eppendorf tube submerged in liquid nitrogen. Müller’s organs from 24 locusts (48 Müller’s organ total) were used for each sample so that we had four samples from young females, four samples from young males, four samples from stressed females, four samples from stressed males. Once all Müller’s organs were on the pestle in the Eppendorf tube Müller’s organs were ground with the pestle for 1 min in the liquid nitrogen and then 2 min at room temperature until the sample started to melt and become see-through. At this point 10 µl of TRizol was added to the sample and then it was ground for a further 3 mins before another 490 µl of Trizol was added. The sample was left on wet ice for 15 mins before 100 µl chloroform was added to the sample, which was then vortxed and left for a further 3 mins on wet ice. The sample was then centrifuged at 12,000g for 15 mins at 4°C and the supernatant (clear top layer) was pipetted into a fresh Eppendorf tube with 250 µl isopropanol and 1 µl glycogen and incubated on wet ice for 10 mins. The sample was then centrifuged at 12,000g for 10 mins at 4 °C. The liquid in the Eppendorf was removed to reveal a pellet. A DNase solution of (45 µl water, 1 µl DNase, 4 µl DNase buffer and 1 µl RNase inhibitor) was used to resuspend the pellet. The Eppendorf was then incubated at 37°C for 15 mins. Ice cold isopropanol 60 µl was added and then the sample was incubated for 10 mins before being centrifuged at 12,000g for 10 mins at 4°C. The liquid was removed and the pellet was dislodged with 70% ethanol before being centrifuged at 7500g for 5 mins at 4°C. The 70% ethanol was removed and the pellet was airdried for 1 min before being eluted in 20 µl DEPC-treated water.

### Sequencing alignment, gene identification, GO analysis

Total RNA samples were sequenced on the Illumina platform by Novogene sequencing at a depth of 60 million reads with 150 base-paired ends (Cambridge). Quality trimmed samples read quality was checked using FastQC (v0.11.5) (Andrews, 2010). Transcriptomic reads were mapped to the iqSchGreg1.2 reference genome (NCBI) using the STAR 2-pass method (v2.7.9a) (Dobin and Gingeras, 2015). Gene count number was generated using HTSeq (Anders et al., 2015). Sample normalization and differential expression analysis were performed using DESeq2 (v1.28.1) (Love et al., 2014) on Rstudio (4.2.1) using the model: design = ∼ condition (with condition: young female, young male, stressed female, and stressed male). To calculate p-values, we used DESeq2’s inbuilt Wald test and p-values were adjusted for multiple hypothesis testing using DESeq2’s inbuilt Benjamini and Hochberg false discovery rate method. Statistically significant genes were those with p-adj <0.05. We did not set a fold change as a threshold. To determine differentially expressed sex-independent genes after stress, we removed all stress induced differentially expressed genes shared between both male and female datasets. Gene ontology (GO) terms were generated using topGO (Alexa and Rahnenfuhrer, 2022) with a false discovery rate <0.05. When listing our top 10 GO terms (Fig. 4), only terms which contained more than 1 gene were included. Analysed datasets are freely available at Medeley Data: doi: 10.17632/4sxj4t7vr6.1

### Statistical analysis

Throughout the manuscript n refers to the number of recorded neurons and N refers to the number of Müller’s Organ preparations used to achieve these recordings (i.e. n = 12, N = 8 means that 12 neurons were recorded from 8 Müller’s Organs). All n numbers are displayed on the figures for clarity. The spread of the data is indicated by 1 standard deviation. Due to stark differences in the sex and the age of the locusts it was not possible to blind the experimenter when collecting the data. This data was, however, was originally collected for an study on auditory decline (Blockley et al., 2022) and we only analysed the data later to find sex differences. All data either remained blinded or was recoded to be completely blind when analysing the data to avoid unconscious bias. To test for differences and interactions between control, and stressed locusts we used either a linear model (LM) or Linear Mixed Effects Model (LMEM), with condition (young or stressed) and sex as fixed effects, and Locust identity and SPL as a random intercept, when repeated measurements are reported. Models were fitted in R (Version 3.4.3) with the package LME4 (Bates et al., 2015). The test statistic for these analyses (t) are reported with the degrees of freedom (in subscript) and p value, which are approximated using Satterthwaite equation (lmerTest package) (Kuznetsova et al., 2017). We report Cohen’s d effect size for significant differences. Curves where fitted to the data using the drm package in R for patch-clamp recordings (Ritz, 2016). The drm package was also used to compute t and p values when comparing young and stressed four-part Log-Linear models. F statistics of the Log-Linear model fits were computed by excluding treatment (young or stressed) as a factor. Higher F statistics donate a stronger effect of treatment. In order to compare responses between young and stressed locusts across SPLs we adopted an approach first implemented in pharmacology research. In our work the ‘‘dose-response curves’’ are equivalent to SPL-transduction current curves. This allowed us to maximise the information contained in each dataset and to quantitatively compare model parameters such as: Hill coefficient (steepness of slope), maximal asymptote (maximum s ratio), and inflexion point (s ratio at the steepest part of the slope). We did this using the drm function of drc package (Version 3.1-1, Ritz et al., 2015). We fitted four-part Log-Linear models with transduction current as the dependent variable with condition (young or stressed) and SPL as the independent variables. We discounted two loglinear fits from two recordings (out of 32) as the loglinear fit gave unrealistic values for the parameters, that would drastically skew the statistical analysis. For model comparisons the standard error is given (as opposed to standard deviation). Analysed datasets and the code to plot and statistically analyse the data are freely available at Mendeley Data: doi: 10.17632/4sxj4t7vr6.1

## Results

### Müller’s organ anatomy and function

We wished to establish if there were sex differences in addition to already known age- and noise-related auditory decline in desert locusts (Warren et al., 2020; Blockley et al., 2022). Each desert locust has two Müller’s organs attached directly onto two tympani in the lateral part of each 1^st^ abdominal segment. The tympanum has a tonotopic travelling wave and ∼80 auditory neurons that attach onto distinct parts of the tympanum to detect a range of frequencies from 200 Hz – 25 kHz (**Figure 1C**). We choose to target Group III auditory neurons as these compose the majority ∼46 out of 80 of auditory neurons in Müller’s organ (Jacobs et a., 1999) and are accessible for single cell electrophysiological recordings (Warren and Matheson, 2018). Hearing thresholds of locusts are as low as 40 dB SPL (Sound Pressure Level) (Michelsen, 1968) and saturate around 110 dB SPL (Warren et al., 2020).

### Whole-cell patch-clamp recordings of the transduction current

We measured sound-evoked transduction currents in individual auditory neurons from Müller’s organs from excised ears (**Figure. 2A**). We isolated and optimized the transduction current at the distal ciliated end of the auditory neuron using pharmacology, voltage protocols, and the optimal sound stimulus (detailed in methods) (**Figure 2B**). For the young group we measured transduction current amplitudes in the 12 hours following mock deafening (Blockley et al., 2022). The transduction current increased in magnitude with an increase in sound amplitude and followed a log linear relationship, with a lower and upper asymptote (**Figure 2C**). We fitted a four-part log-linear model to the transduction currents as a function of the sound pressure level for the averaged data for males and females of the young cohort (**Figure 2C**, solid lines). For statistical analysis we fitted the four-part log-linear model to transduction currents from averaged individual locusts. We ran a linear model on the three model parameters and found no sex differences for the young cohort in: hill coefficient (t_(29)_=0.088, p=0.93), maximum asymptote (transduction current) (t_(29)_=0.525, p=0.60), inflection point (t_(29)_=-0.119, p=0.91). We also found no differences in the magnitude of the channel opening events (t_(31)_=-0.119, p=0.91). We measured the electrophysiological properties of the auditory neurons. There was no difference in the membrane resistance ((t_(31)_=0.892, p=0.379) and capacitance (t_(31)_=-1.29, p=0.21) but there was a difference in the resting membrane potential (t_(31)_=2.14, p0.04, Cohen’s d = 0.76) with male auditory neurons 7.1 ± 3.3mV more depolarised than female. To quantify the difference in the log-linear fit, utilising all the data, we compared models of the average transduction currents which gave an F statistic of 5.23.

**Figure 2.**
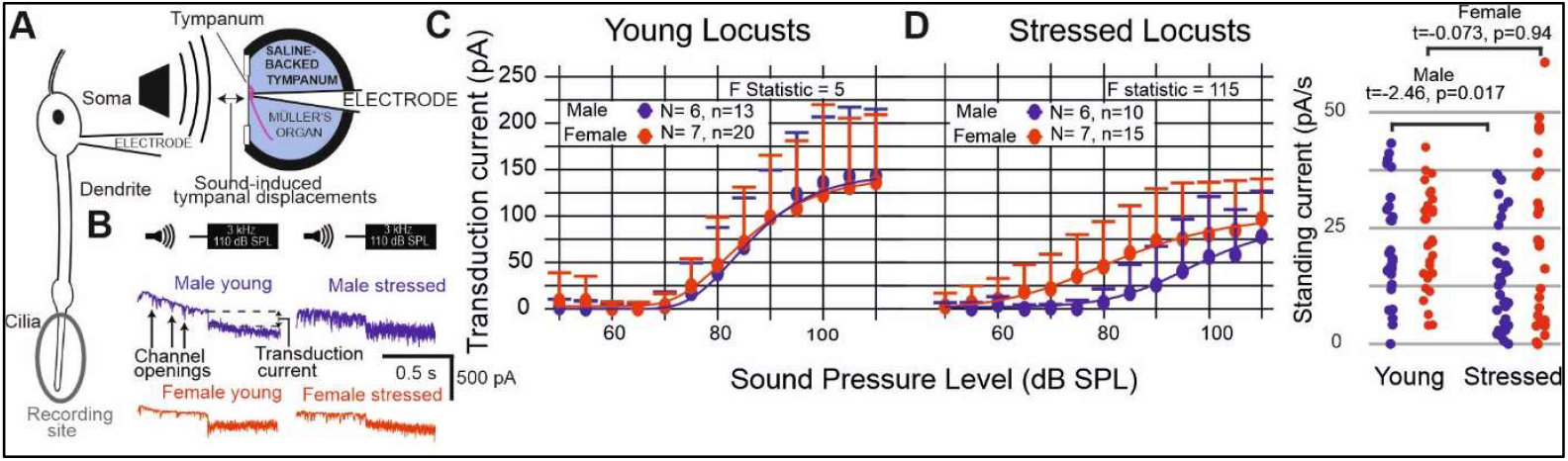
Transduction current recordings for female and male auditory neurons for young and stressed locusts. **A**. Experimental setup showing configuration for whole-cell patch-clamp recordings. **B**. Example voltage-clamp recordings of the transduction current. The neuron is clamped at -100 mV to increase the driving force for the transduction current and a 3 kHz tone is played at 110 dB SPL. **C**. Transduction currents measured from the auditory neurons of young female and male locusts. Dots donate the average, error bars are the positive standard deviation and the solid lines are Log-linear models fits to the average data. **D**. Transduction currents measured from the auditory neurons of old female and male locusts exposed six times to noise. Dots donate the average, error bars are the positive standard deviation and the solid lines are Log-linear models fits to the average data. **E**. Comparison of the standing current in young and stressed Müller’s organs split by sex.

We compared log-linear model fits of the transduction current for the young and stressed cohort (independently of sex) and found no difference in the hill coefficient (t_(57)_=-1.118, p=0.27), inflection point (t_(57)_=1.62, p0.11), maximum transduction current (t_(57)_=-1.12, p=0.267). However, when we computed the maximum transduction based on the log-linear model we found a strong trend for a lower transduction current of 59 ± 15 pA at the maximum SPL (t_(57)_=-3.84, p=0.0003, Cohen’s d=0.99). There was a significant reduction in the amplitude of channel openings (t_(59)_=-3.34, p=0.001, Cohen’s d=0.84) but other electrophysiological properties of auditory neurons were unchanged between young and stressed locusts: membrane resistance (t_(59)_=0.329, p=0.74), capacitance (t_(59)_=-0.07, p=0.94), and resting potential (t_(59)_=1.169, p=0.25).

Next, we focused on sex differences in the stressed cohort. We found no difference in the hill coefficient (t_(25)_=2.11, p=0.83), but did find a difference in the inflection point (t_(25)_=2.14, p=0.04, Cohen’s d=0.84), with male’s inflection point shifted 15 ± 6.7 dB SPL higher (**Figure 2D**). The maximum transduction current was not different (t_(25)_=-0.417, p=0.68) when comparing the log-linear fit and even when calculating transduction current amplitudes at 110 dB SPL from the log-linear model (t_(25)_=-0.936, p=0.358). However, in the stressed cohort the difference between the log-Linear model fit of the male and female data significantly increased (F statistic = 184.4). There was a trend for smaller amplitude channel openings for males (t_(25)_=-1.94, p=0.06). We also measured the standing current (in the absence of auditory stimulation) flowing from the neurons cilia, housing the transduction channels and surrounded by receptor lymph, to the soma, where the recording electrode is located. This is an indirect measure of the metabolic activity of the scolopale cell (**Figure 1B**) and the neuron. We found a significant decrease for the standing current for males but not females (males: t_(59)_=-2.46, p=0.017, Cohen’s d=0.35; females t_(54)_=-0.073, p=0.94 (**Figure 2E**). All other electrophysiological properties of the auditory neurons were not different: membrane resistance (t_(25)_=-0.07, p=0.94), capacitance (t_(25)_=0.235, p=0.82), and resting potential (t_(25)_=-1.49, p=0.15).

### Gene expression of Müller’s organ in young and stressed locusts

To understand the physiological process through which the Müller’s organs of females retains better function we extracted RNA from the Müller’s organs of young and stressed male and female locusts. First, we identified differentially expressed genes between male and female Müller’s organs in the young and the stressed cohorts (Supplementary Table 1). We found 39 genes differentially expressed between males and females, with 39 genes differentially expressed in the young cohort and only 9 differentially expressed in the stressed cohort (**Figure 3**). The most highly differentially expressed gene is juvenile hormone (JH) but within the top 12 expressed genes are vitellogenin, greglin, phenoloxidases, prisilkin-39-like, and double sex. The only identified gene upregulated in young males is a probable cytochrome 450.

**Figure 3.**
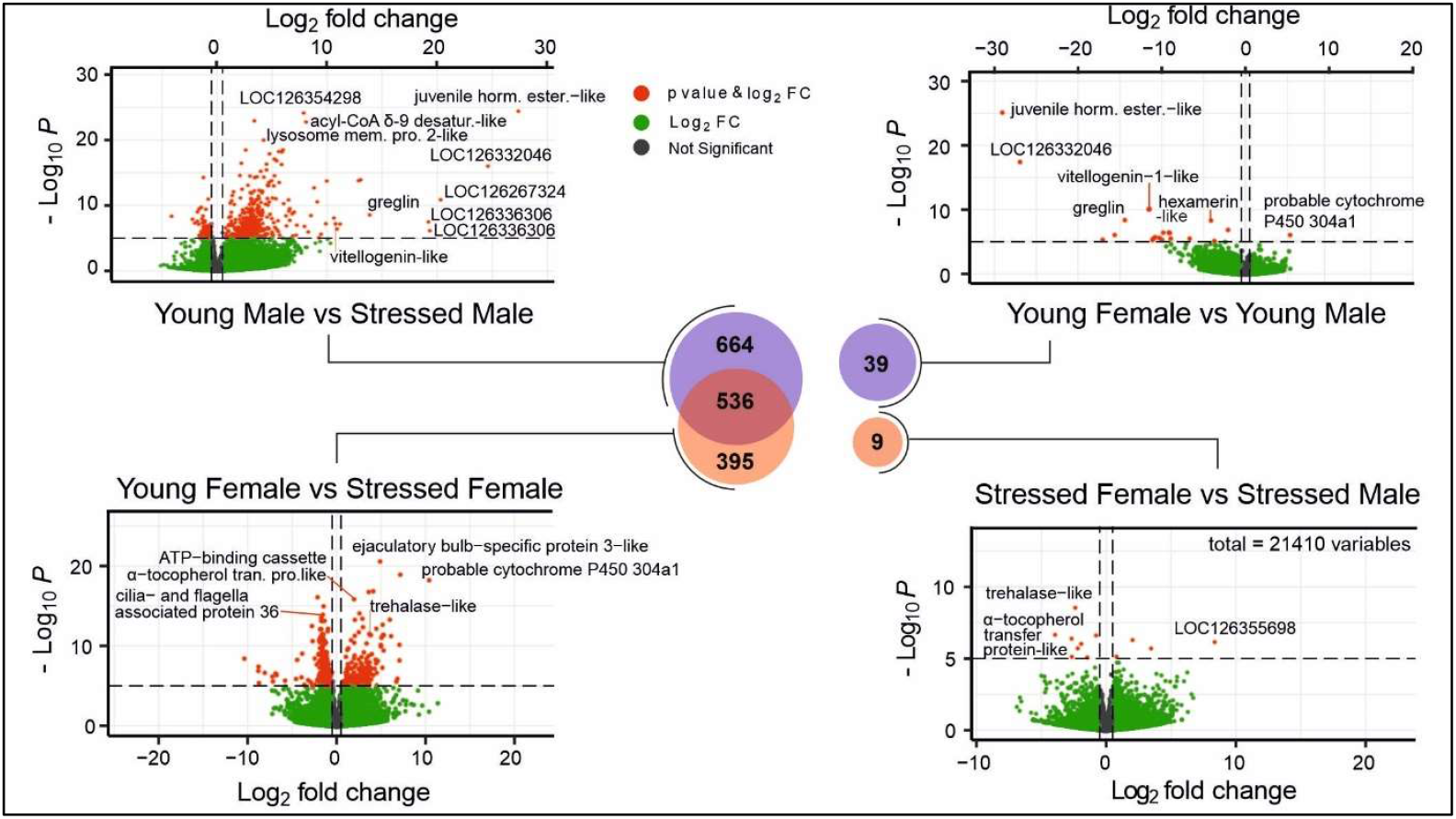
Differential expression Venn Diagram and Volcano plots for the four comparisons: Young Male vs Stressed Male; Young Female vs Stressed Female; Young Female vs Young Male; Stressed Female vs Stressed Male. Venn diagram numbers indicate number of differentially expressed genes. Only genes marked in red on volcano plots sufficiently meet our criteria of fold change and p-value for differential expression. The most highly differentially expressed genes are labelled on the volcano plots.

Through gene ontology analysis we found biological processes and classified them into three categories: immune-related, hormonal-related and neurotransmitter related (**Figure 4**). The top two GO terms upregulated in young females, compared to young males are melanin-associated processes: melanin biosynthetic processes and melanin encapsulation of foreign particles (**Figure 4A**). Within the top 12 are dopamine and ammonium metabolic processes, immunity related: defence response to bacteria and fungus, and then two for analia (most posterior appendage of insects) development. In the stressed cohort, only 4 GO terms were generated, due to the low number of genes in this comparison (**Figure 4B**).

**Figure 4.**
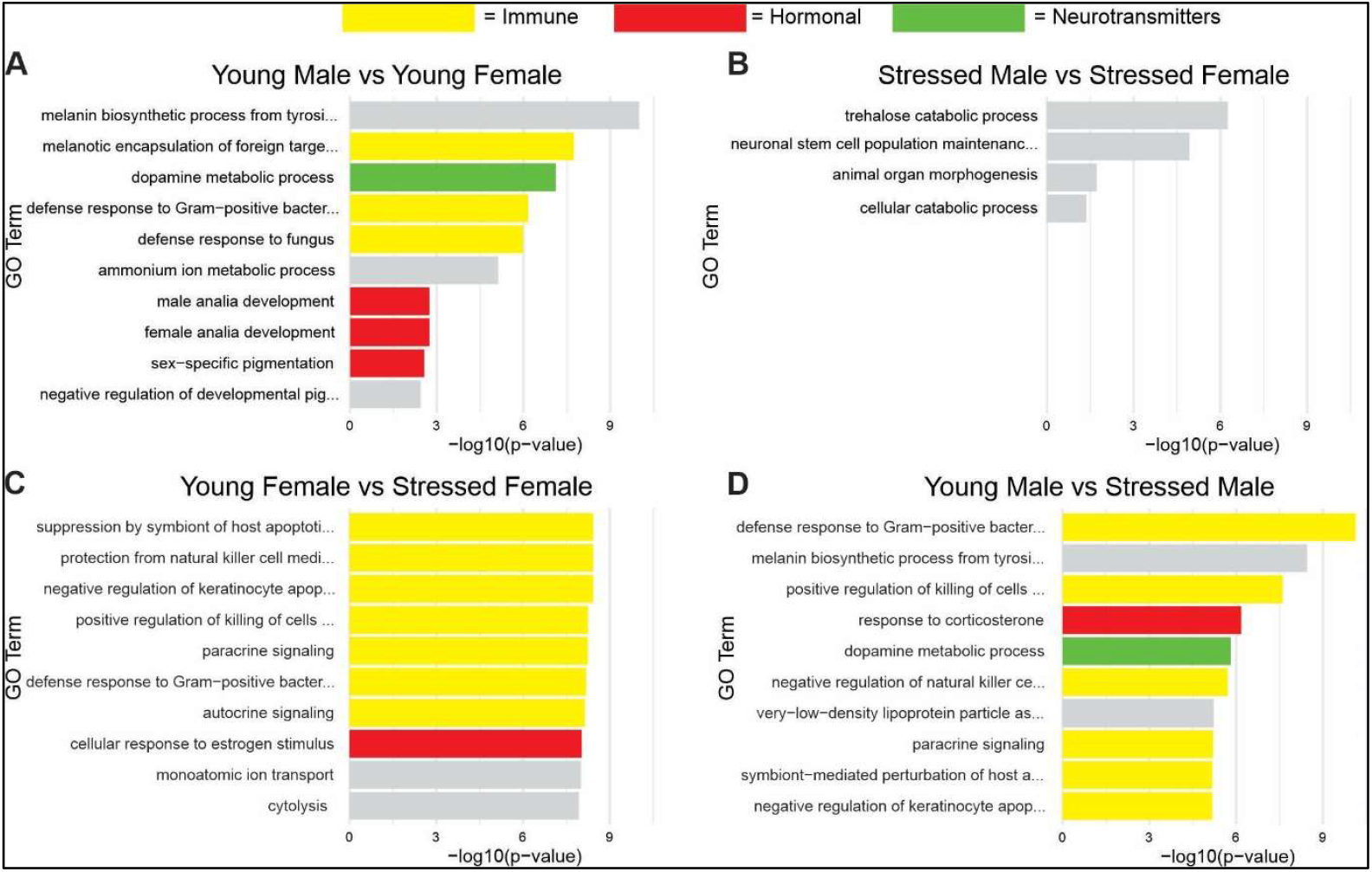
Top 10 GO terms for each comparison in Figure 3. For the two sex-independent analyses, differentially expressed genes contained within both Young Female vs Stressed Female and Young Male vs Stressed Male (the overlap of the Venn diagram in Fig. 3) were removed before carrying out the analysis.

The number of genes differentially expressed between young and stressed female Müller’s organ was 931 and between young and stressed male locusts was 1200 (**Figure 4 C, D**). Of these 536 genes were shared between male and female Müller’s organs. We would therefore expect these genes, (and thus the GO terms they contribute towards) to represent processes not critical in sex differences in Müller’s organ function. Here, very few of the GO terms were cuticle- or immune-related or neural-related, three broad categories used to classify differences in young and aged Müller’s organs (Austin et al., 2024). We found juvenile hormone acid O-methyltransferase-like that generates the active form of JH and the neuropeptide intermediate between corazonin and adipokinetic, adipokinetic hormone/corazonin-related peptide receptor variant I-like.

We analysed genes differentially expressed between young and stressed males (666) and between young and stressed females (395) that were not shared between the sexes i.e. male-specific and female-specific young vs stressed genes. We used GO analysis of male-specific and female-specific young vs stressed Müller’s organs, finding fewer GO terms in the female comparison, the top 10 of which had a considerably lower p-value (**Figure 4**). For males we found the GO term androgen metabolic process, which includes male-specific hormones related to sexual differentiation, prostaglandin (hormone-like compounds) metabolic processes and dopamine metabolic processes for males. In the top 50 GO terms we found only five immune-related GO terms. In females we found three GO terms linked to estrogen: estrogen catabolic process, cellular response to estrogen stimulus, cellular response to estradiol stimulus. We also found 20 immune-related GO terms and five protein modification GO terms. Amongst the top 10 GO terms were 5 GO terms related to glucuronidation of bilirubin.

## Discussion

### Female locusts have better auditory function under stressed conditions

We wanted to investigate if sex differences exist in auditory function in the Desert Locust. If sex differences exist, we wanted to identify the sex with better auditory function and then use gene expression analysis to understand its physiological basis. We found sex differences in auditory function of the Locust’s auditory Müller’s organ, with females retaining better auditory function, but only for older locusts subjected to repeated deafening and aging paralleling that found for older humans (Corso, 1963; Pearson et al., 1995; Homans et al., 2017). Humans, in these studies, were not exposed to noise in controlled experimental context but increased noise levels endemic in industrialised societies are thought to be a contributor for hearing loss (Rosen, 1962). Sex differences were also found in young mice after exposure to damaging levels of noise with better auditory brainstem responses (ABRs) in females, (Milon et al., 2018). Better or more resilient auditory function also appears to hold true for female zebrafish (Han et al., 2022). We also found that the standing current, a measure of the metabolic work of the scolopale cell to maintain the receptor lymph, reduced for males but was maintained for females. This suggests that female’s metabolism is better maintained under stressed conditions. In mammals the stria vascularis serves an equivalent role as the scolopale cell; it maintains the endocochlear potential through actively transporting ions up their electrochemical gradient and is thought to be one of the contributors to noise-induced (Yamane et al., 1995) and age-related hearing loss (Lang et al., 2023; Schuknecht & Gacek, 1993).

### Sex differences in gene expression are more numerous in young Müller’s organs

In young locusts with no differences in auditory function we found 39 genes differentially expressed between the sexes with 37 upregulated in females and only two more highly expressed in males. The most highly differentially expressed gene is juvenile hormone (JH) and then half of the next 12 top expressed genes are vitellogenin and greglin, which are proteins, with a wide range of functions, under the control of JH. Vitellogenins are produced by the ovary and fat bodies and have diverse roles in homeostasis, protein synthesis and immunity. Also upregulated in female Müller’s organs are invertebrate immunity-related genes such as phenoloxidases, a cuticle (chitin) regulation gene, prisilkin-39-like, and double sex, involved in sexual differentiation. One gene, a probable cytochrome 450 is higher in male Müller’s organs and is involved in the synthesis and degradation of steroids and JH and the metabolism of foreign molecules. Thus, a natural conclusion is that the upregulation of JH in female Müller’s organ and its associated proteins is due to a downregulation of JH in males. In other insects, such as the cockroach JH is higher in females and confers a greater resistance to both biotic and abiotic stressors such as bacteria and hydrogen peroxide (Liu et al., 2024). We have to also accept that higher JH expression, and the proteins it stimulates, in female Müller’s organ is unrelated to differences in physiological function of the auditory system. JH is important for egg maturation (Maeno & Tanaka, 2009) and expression in Müller’s organ and could be part of a generic expression programme in a wide range of tissue types in females. All of these gene changes are maximal in young male and female locusts when there is no difference in auditory function. We identified melanin as the top two upregulated biological process in female Müller’s organ, followed by dopamine and immunity-related GO terms. Interestingly the racial differences in human hearing loss, where black ethnicities have less hearing loss than their white counterparts (Lin et al., 2011), are hypothesised to be due to the higher expression of melanin in the stria vascularis which acts as a free-radical scavenger stemming free-radical damage (Murillo-Cuesta et al., 2010). Perhaps this mechanism has been exploited by female Müller’s organ that also have melanocytes (Gray, 1960).

### The majority of sex differences in genes manifest during aging and noise-exposure

Locusts exhibit age-related and noise-induced reduction in auditory function (Warren et al., 2020; Blockley et al., 2022) but the majority of gene expression changes are due to aging (Austin et al., 2024). It is perplexing to find that differential gene expression between the sexes reveals only 39 genes and 9 genes in young and stressed Müller’s organs respectively. We found that an overwhelming majority of the differentially expressed genes were through comparisons of young and stressed Müller’s organs, with about half of the genes shared between the sexes (536/1200 for males and 536/931 for females). This makes 664 male-specific Müller’s organ genes, and 395 female-specific Müller’s organ genes, differentially expressed under stressed conditions. If sex differences in auditory function are controlled by hormones and neuromodulators, the steroid estrogen would be a clear front-runner for females as we found three estrogen-related GO terms in the top 50 differentially expressed genes. Here, estrogen GO terms likely reflect estradiol and estrogen-related receptors (ERRs). In mammals there is a clear president for estrogen’s role in limiting auditory decline. A single nucleotide polymorphism in the estrogen-related receptor gamma gene is associated with hearing in human females only (Nolan et al., 2013), its absence results in increased hearing thresholds in mice (Simonoska et al., 2009; Nolan et al., 2013) and its replacement in rats after ovariectomy reduced noise-induced threshold shifts (Kim et al., 2020). Although sex determination in insects was thought to be solely genetically based (Mechoulam et al., 1984) there are clear sex-specific hormones expressed in insects (De Loof & Huybrechts, 1998) and the migratory locust expresses the steroid estradiol (Novak & Lambert, 1989). ERRs are ancient and existed in the ancestor of bilaterian metazoans and insects still retain ERRs orthologous with, and with the typical structural features of, mammalian ERRs (Bardet et al., 2006). The role of ERRs in insects is diverse and not broadly studied. In flies EERs are involved in: modulation of carbohydrate metabolism genes, axoneme assembly, mitochondrial function, and a key metabolic transition (Kovalenko et al., 2019; Misra et al., 2017; Tennessen et al., 2011). In bees ERRs are key for responses of larvae to different nutritional environments (Santos et al., 2016) and in moths ERRs are necessary for pheromone responses (Bozzolan et al., 2011). ERRs seem to be broadly implicated in metabolic processes and could be directly involved in mitochondrial regulation if insects, like mammals, process ERRs in mitochondria (Yang et al., 2004). We found a decrease in the standing current for male Müller’s organs indicative of decreased metabolism in males compared to females. We postulate that these metabolic signatures are key to the sex-differences in auditory function that emerge under stressed conditions. Possible mechanisms are already identified in mice; estrogen interacts with the mammalian equivalent of the insect’s scolopale cell (marginal cells in the scala media) to regulate ion flow (Lee & Marcus, 2001). Sex differences in metabolism are well established both in insects (Shingleton & Vea, 2023) and humans (Kastenmüller et al., 2015) and could explain sex differences in auditory decline across diverse animal phyla.

For male-specific GO terms differentially expressed between young and stressed Müller’s organ we find positive regulation of mitophagy and in the female equivalent we find negative regulation of mitochondrial DNA replication. This hints that the mitochondria in stressed males fare less well than female mitochondria. Female-specific GO terms between young and stressed Müller’s organs also reflect higher regulation of metabolic processes with 9 out of the top 50 GO terms compared to the male’s 5 out of 50. The fewer genes and less GO terms overall that change within females with stress (compared to within males) also suggests a more robust system in females that is less susceptible to transcriptional dysregulation (or attempted repair) with age and noise-exposure. The majority of differentially expressed genes in both males and females were found to be downregulated. There are 171 upregulated genes vs 224 downregulated in females, and in males, this difference is even more exaggerated, with only 16 genes upregulated, and 650 genes downregulated with age and noise-exposure.

We found broader hormonal GO terms for androgens and prostaglandin (hormone-like compounds) metabolic process in males, which were differentially expressed between young and stressed. We previously found that genes differentially expressed between young and old male Müller’s organs and Müller’s organs exposed to noise fell into three broad categories: cuticle-related, immune-related and neural (Austin et al., 2024). We again found many immune-related GO terms but females had a larger number of immune-related (20) genes compared to males (5). There were a further 14 immune-related GO terms shared between male and female Müller’s organs, suggesting that upregulation of immune function is common in both sexes.

## Conclusion

We found that females retained better auditory function when aged and exposed to repeated bouts of noise exposure (stress). Most the genes differentially expressed between males and females manifested during aging and over repeated noise exposures. We suggest that auditory resilience in females is established through the expression of JH and female-specific steroids like estradiol that have downstream affects, such as protein modifications and repair, better immune responses that protect Müller’s organ and better regulation of metabolism. We found that female Müller’s organs maintain better metabolic function in stressed conditions. We speculate that better metabolism through the action of sex-specific steroids on mitochondria is the most likely explanation for the sex differences in auditory function of the desert locust.

